# Prior EEG marks focused and mind-wandering mental states across trials

**DOI:** 10.1101/2021.09.04.458977

**Authors:** Chie Nakatani, Hannah Bernhard, Cees van Leeuwen

**Affiliations:** Brain and Cognition Research Unit, KU Leuven, Tiensestraat 102 - Box 3711, 3000 Leuven, Belgium; Department of Cognitive Neuroscience, Faculty of Psychology and Neuroscience, Maastricht University, Maastricht, The Netherlands; Center for Cognitive Science, RPTU Kaiserslautern, Kaiserslautern, Germany

**Keywords:** Spontaneous mental process, dynamical modeling, EEG alpha amplitude, event related potential, cognitive control

## Abstract

Whether spontaneous or induced by a tedious task, the transition from a focused mental state to mind wandering is a complex one, possibly involving adjacent mental states and extending over minutes or even hours. This complexity cannot be captured by relying solely on subjective reports of mind wandering. To characterize the transition in a mind-wandering-inducing tone counting task, in addition we collected subjective reports of thought generation along with task performance as a measure of cognitive control and EEG measures, namely auditory probe evoked potentials (AEP) and ongoing 8-12Hz alpha-band amplitude. We analyzed the cross-correlations between timeseries of these observations to reveal their contributions over time to the occurrence of task-focused and mind-wandering states. Thought generation and cognitive control showed overall a yoked dynamics, in which thought production increased when cognitive control decreased. Prior to mind wandering however, they became decoupled after transient increases in cognitive control-related alpha amplitude. The decoupling allows transitory mental states beyond the unidimensional focused/wandering continuum. Time lags of these effects were on the order of several minutes, with 4-10 minutes for that of alpha amplitude. We discuss the implications for mind wandering and related mental states, and for mind-wandering prediction applications.

## Introduction

Studies in mind wandering typically involve contrasting self-reported task-focused and mind-wandering states. However, the transitions between these states can last minutes or even hours in wakeful individuals, and may be too complex for a simple dichotomy. A continuum of task-focused and mind-wandering states would be insufficient either. The mental states visited in the transition may not fall on a unidimensional continuum (Mittner et al., 2016; Nakatani et al., 2019). We aim to characterize these transitions using a broader set of behavioral indicators in combination with electrophysiological measurement, and apply cross-correlation to determine the time course of their effects, and their directionality, on mind wandering behavior.

One candidate indicator will be the rate of thought generation. The wakeful mind spontaneously generates thoughts including those that have no immediate relevance to the current situation. Every thought that occupies the mind offers an opportunity for distraction. “Having multiple thoughts”, therefore, is potentially a factor predicting mind wandering. Introspection and experience sampling studies have asked participants to identify multiple distinct thoughts that occupy their mind at a given time and report their number (Campbell, 1960; Ganschow, 2017; Nakatani et al., 2019). Even though some mental content may not be consciously accessible (Dijksterhuis & Nordgren, 2006), subjectively reported numbers of thoughts were found to correlate positively with the propensity of mind wandering (Ganschow, 2017). Their reported number, therefore, can be considered a measure of the mental state relevant to mind wandering.

Multiple thoughts on one’s mind as such is not a sufficient condition for mind wandering. It is possible, at least in principle, that all these thoughts are task-relevant. Allowing the mind to wander away from task-relevant thoughts may suggest perturbation of the intentional and goal-directed behavioral dispositions collectively known as cognitive control (Cohen et al., 1990; Funes et al., 2010; Otto et al., 2015; Ridderinkhof, 2002). An episode of mind wandering could occur as cognitive control slips away in a boring and repetitive task (Smallwood & Schooler, 2006). Still, certain control functions may be retained, in particular over the operation of working memory (McVay & Kane, 2012), for monitoring contextual factors of mind-wandering such as task difficulty (Levinson et al., 2012), and even for willfully deciding to initiate mind wandering (Seli et al., 2016). Either way, whether cognitive control slips away, for instance due to fatigue after some time on task, or when mental resources are deliberately reallocated to task-irrelevant processes, task performance will suffer. Measures of task performance will thus reflect the availability of cognitive control for the task.

Given the number of thoughts and task performance as behavioral measures of mind wandering, we expect it likely to occur when the number of thoughts is high and task performance is poor. Having adopted a dynamic approach to mental states (Mittner et al., 2016), changes over time in the number of thoughts and a measure of visual task performance in an experience sampling stud helped identify transitions between focused and mind wandering states (Nakatani et al., 2019). Spontaneous, unintentional mind wandering over the wakeful hours of the day was driven by an increase in thought production in combination with a decrease in cognitive control. A dynamic approach situates mind-wandering within a space of “spontaneous thought” states (Christoff et al., 2016), where a low number of thoughts combined with poor performance of the primary task, for instance, might suggests rumination or obsessive thinking; a high number of thoughts together with a high task performance might characterize multi-tasking, creative thinking, or even an intentional mind wandering state.

Mapping out the transition from a focused to a mind wandering state showed it to occur most often by passing through a “multi-tasking” state (Nakatani et al., 2019). This observation flouts the notion that thought generation and cognitive control are yoked in a one-dimensional dynamic. Such a dynamic would seesaw the rate of thought generation and the primary task performance (if one goes up, the other goes down). A state where thought generation is high *and* task performance is high would never be reached. Therefore, the transition via this state suggests that here at least, our two measures might represent independent contributions to the dynamics of mental states.

Let us assume that both our measures reflect autonomous changes over time in, respectively, thought generation and cognitive control. Sooner or later, these changes will manifest themselves in self-reported mind wandering. Our aim is to determine the timing (how much sooner or later) and direction (whether positive or negative) changes in our measures are reflected in behavior.

To find the critical time, or times, at which these changes set the mind to wander, we record brain signals. Since timing is our emphasis, our signal of choice is EEG. Another reason for using EEG is that the current study should have potential applications outside of the laboratory. The downside of EEG is its poor spatial resolution, and the resulting inherent uncertainty in spatial mapping between brain signals and high-level mental functions (Irving & Thompson, 2018; Metzinger, 2018). For this reason, we rejected using spatial mapping for selecting the relevant signals and instead chose ones with high temporal resolution, suitable for pursuing stochastic causality (Granger, 1969). In this perspective, if a given brain signal X explains the system’s output Y (e.g., the number of thoughts) more than its own history of Y (e.g., the past number of thoughts), then X could be considered as an effective component of the system. Such an effect could be observed for specific time lags (latencies) between X and Y.

Two EEG measures were adopted. One measure is amplitude of the auditory evoked potential (AEP) to tone stimuli, which is closely linked to auditory signal processing (Picton & Hillyard, 1974; Picton, Hillyard, Krausz, & Galambos, 1974), and the other is amplitude of alpha-band (8-12Hz) activity, which constitutes a measure of on-going brain activity sensitive to changes in mental state, e.g., between resting and task states (Kam et al., 2022; Lehtelä et al., 1997; Tiihonen et al., 1991; Weisz et al., 2011).

The timescale for the cross-correlation analysis was determined considering studies relevant to the deployment of cognitive control and thought generation. Sustained attention studies have observed lapses of attention on the timescale of milliseconds and seconds (Cheyne et al., 2006; Funes et al., 2010; Redmond G. O’Connell et al., 2009). Vigilance studies have reported loss and recovery of cognitive control on the timescale of hours in the range of ultradian rhythm (1.5-several hours/cycle). Similarly, the rate of thought generation changed in the ultradian range (Nakatani et al., 2019). These studies did not address how such loss of control leads to an episode of mind wandering. In addition, the timescales in the previous studies are either too fine (ms, s) or too coarse (hours) to monitor an episode of mind wandering. In an experience sampling study of mind wandering, an average duration of mind wandering of 10 min was observed (Ganschow, 2017). To capture the transition to mind wandering over such a period, our current analyses used a timescale of minutes.

Data were collected in an experiment in which participants counted tones presented in a slow sequence. Subjective reports of mind-wandering were collected after every trial (i.e. approximately every 2 min), in order to capture the 1^st^-person perspective report of mind wandering (Metzinger, 2018). In addition, subjective reports of the number of thoughts were collected as a measure of the spontaneous thought generation rate, while the percentage of correct tone count was used as the measure of the availability of cognitive control for the task. EEG was continuously recorded during the experiment. We performed cross-correlation on the timeseries of our behavioral and EEG measures, to which a multivariate general linear model was applied. The results suggest that thought generation and cognitive control have distinctive contributions to mind wandering. Their seesaw effect is disrupted by an effect of alpha activity on cognitive control. Alpha activity affected cognitive control 4 to 10 minutes before mind wandering was reported. We discuss the implications of our results for understanding the mental dynamics for mind wandering, along with possible applications such as predicting upcoming episodes of mind wandering.

## Methods

### Overview

Participants performed a task of counting tones up to a target number (between 20 and 24) pre-specified at the start of each trial. Each time after reaching the target, they retrospectively rated their mental states during the trial on a scale, ranging from fully task focused to fully mind wandering (%MW), and provided a subjective report of the number of thoughts (NoTs) that had occurred during the trial (Figure 1, top). NoTs were used as measure of thought generation rate (Nakatani et al., 2019), while performance of the tone count, namely, the percentage of correct count (%CC) was used as measure of cognitive control available for the task. EEG was recorded continuously during the counting task, in order to compute amplitude of the auditory evoked potential (AEP) to the tone stimuli as measure of sensory information processing and amplitude of alpha-band (8-12Hz) activity as measure of ongoing brain activity. (See **Computation of AEP amplitude** and **Computation of alpha-band amplitude**)

**Figure 1.**
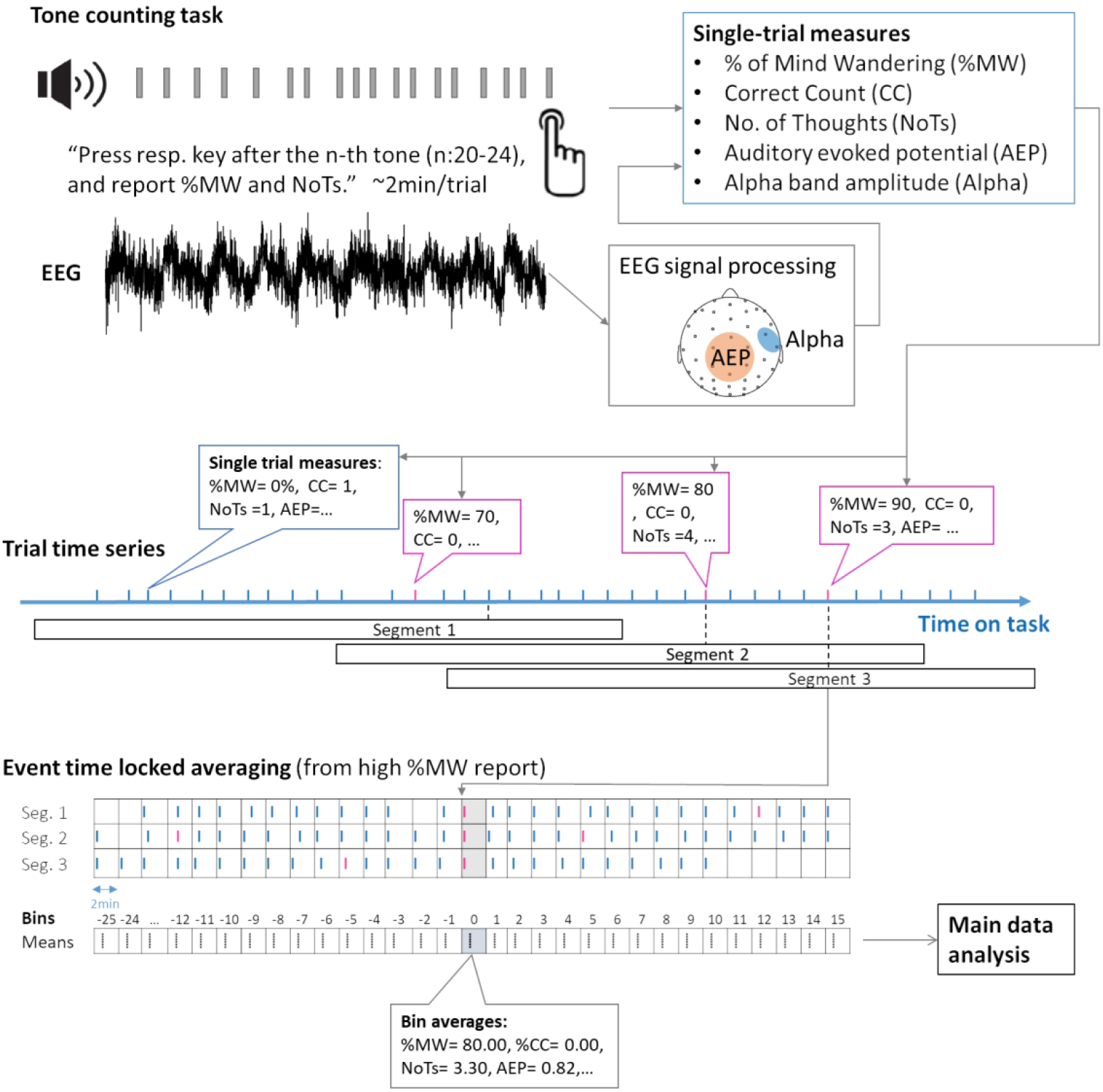
Tone counting task and event time-locked averaging. Tone counting task: Participants were asked to count n (20-24) tones and press a key when the count was reached. After the key press, they reported the percentage of time they spent on mind wandering (%MW) and the number of thoughts (NoTs) during the trial. EEG was recorded during the task. Amplitude of alpha activity and auditory evoked response (AEP) were computed from the EEG. Each trial thus yields 5 measures, Correctness of the count (CC), NoTs, %MW, Alpha, and AEP amplitude. On NoTs, %CC, Alpha, and AEP amplitude, event-time locked averaging was performed separately from the moments of mind- wandering (%MW≧70) and task-focused (%MW < 30) reports, respectively. Segmentation is illustrated for mind-wandering reports and was performed likewise for task-focused reports. Segmented timeseries were aligned and averaged. The averaged timeseries were used in the main data analysis.

As reports of mind wandering, we considered instances of reported %MW higher than 70%. Timeseries (technically, interval series) of NoTs, %CC, AEP, and alpha amplitude were segmented and averaged form each time point of a report if mind wandering and averaged over the reports (Figure 1, middle). The event time-locked averaging reduced components which are not time-locked to the corresponding subjective report, similarly to the way ERPs are extracted from averaged signals (Nakatani et al., 2019). Event time-locked averaging was likewise performed from the time points of task-focus reports (reported %MW less than 30%) (See **Preprocessing of subjective reports and behavioral data**, and **Event time-locked averaging of AEP and alpha**.)

Effects of the EEG measures, AEP and Alpha, on the psychological measures, %CC and NoTs, were estimated for the task-focused and mind wandering report conditions separately using multi-variate multiple regression models. The models were designed to identify the neural effects with a latency between 0 and 20min in 2-min. steps. Statistical significance of effects were estimated in terms of stochastic causality (Granger, 1969). (See **Models and main data analysis**.)

The multivariate analysis enables effects of both AEP and Alpha measures on %CC and NoTs to occur independently, and to detect their specific time lags. Previous studies did not include neural effects with latency longer than 1min. We considered effect latencies between 0 and 20min.

### Ethical approval and Participants

The study was approved by The KU Leuven Medical Ethics committee (protocol number: S56128). Twenty healthy volunteers (average 23.00 years, 6 male) participated in exchange for monetary compensation (10€/hour). They were recruited via the online participant recruitment system at KU Leuven. Prior to participation, they were given information about the study and gave oral and written informed consent. All data were collected prior to the onset of the 2020 COVID19 pandemic.

### Procedure

Experiments were conducted in single sessions with individual participants, in an EEG-laboratory of the University Hospital of Leuven. After electrodes were attached, the participant was seated in a dimly lit room in front of speakers and a monitor. Baseline EEG was recorded (eyes open/relax 1 min, and eyes closed/relax, 1 min). This was followed by an auditory oddball task for the estimation of evoked response parameters, and next by the tone counting task. EEG was recorded during both tasks. Afterwards, baseline EEG measurement was performed again. Then the electrodes were removed. The participant were debriefed before leaving. The experimental took about 3 hours in total.

### Tasks

To elicit a sizeable incidence rate of mind wandering during an approx. 2-hour period, a *tone counting* task was used (Figure 1, top). The task consisted of 37 trials lasting approx. 90s each, in which a sequence of brief tones, 100ms in 440Hz frequency, was presented at 70dB via the speakers located at right and left sides of a monitor in front of the participants. The tones were presented with a stimulus onset asynchrony (SOA) between 2000 and 2500ms (pseudo-randomized with 100ms steps). At the beginning of each trial, a number was presented in the center of the display. The number indicated how many tones had to be counted before a response. The number ranged between 20 and 24 (pseudo-randomized over trials). Participants put their right index finger on Key “5” of the ten-key pad and closed their eyes before the series of tones started. When the target number was reached, participants responded by pressing the key. Then they opened their eyes and reported the percentage of time spent on task irrelevant thoughts in 0 to 100% in 10% steps. An eleven-point scale (0% --10%--….-90%--100%) was shown on the display and participants reported the percentage (%MW) using the ten-key pad. Next, participants reported on the keypad the number of thoughts (NoTs) that had occurred in their minds during the trial on a ten-point scale. The scale ranged from 1: only task-related thought to 10: task-related and 9 or more task-irrelevant thoughts. Participants were instructed to evaluate the number of thoughts in a consistent manner over trials. For instance, they were instructed to report “1” when they had only task-related thoughts; when they had one task-irrelevant thought, for instance about having lunch after the experiment, they reported “2”. They were told that items included in the thoughts (e.g., a sandwich, an apple, and a bottle of soda) were not counted separately. If two different thoughts occurred at different times during the trial (e.g., thoughts on lunch, then later during the trial thinking about doing laundry), they reported “3,” and so on. The instruction was the same as that in Nakatani et al., (2019). Participants had the opportunity to take a break at marked sections of the task. EEG was recorded throughout the task.

An *Auditory Oddball Task* was applied prior to the tone-counting task to specify the relevant parameters of the auditory evoked potentials (AEP), namely, scalp distribution, latency, and main frequency band, for subsequent use in the tone counting task. Participants placed their right index finger on the “5” key of the ten-key pad and closed their eyes. Then, a series of 110 tones was presented at 70 dB via the speakers, with an SOA pseudo-randomized between 2000 and 2500ms in 100ms steps. Of these tones, 101 were standard (440Hz, 100ms duration), physically identical to the ones used in the tone counting task, and 5 were odd-ball tones (510Hz, 100ms duration), which were randomly interspersed. Participants were asked to respond to the oddballs by pressing the key as fast as possible.

Both tasks were implemented using PsychoPy Version 1.82.01 (Peirce, 2007) on Windows XP.

### EEG recording

EEG was recorded using the Patientguard EEG recording system (ANT Neuro, Enschede, The Netherlands). Of 64 channels in total, 59 were EEG channels and the remaining ones were used for recording the electrooculogram (EOG, vertical and horizontal) and recording reference (right mastoid, which was later re-referenced to average reference). The placement of the EEG electrodes was according to the manufacturer-defined whole head system rather than the 10-10 system. The signal was sampled at 2048Hz. Electrode impedance was 20 kΩ or less (for the majority of electrodes, the impedance was 5 kΩ or less).

### Preprocessing of subjective reports and behavioral data and event time-locked averaging

Each trial of the tone counting task had 5 measures, NoTs, CC, %MW and two EEG measures (Figure 1, top). The measures are treated as timeseries (Figure 1, middle). Timeseries of NoTs and %CC were segmented for event time-locked averaging, relative to the time of a high- or low %MW report. Thresholds for high and low %MW, 70% and 30%, respectively, were decided based on the grand-average histogram of %MW (Appendix Fig. A1). From the time of the report (Time0), the NoTs and %CC series were segmented between −50min and +32min, allowing overlap between segments. The extension to the past enables estimating autoregressive effects. On average, 9.11 and 6.53 segments per participant were obtained for high- and low %MW reports, respectively. In both types of reports, segments were aligned for averaging from Time0. However, trials do not necessarily align in time (except for those including Time0 of a given high- and low %MW reports), since the inter-trial interval varied according to the number of tones per trial and to individual differences in report timing. Thus, we set bins of 2-min intervals (Figure 1, middle); the data points between Time0 and 1min 59.99s were averaged and considered as the data for Bin0. Similarly, the next interval was between 2min and 3min 59.99s, and the averages were data for Bin1 and so on (Figure 1, bottom). The averaging cancels effects that are not time-locked to high%MW or low%MW reports. Canceling these non-time locked effects by averaging is analogous to removing non-time locked event noise by averaging EEG segments to obtain ERPs. Further details on the event time-locked averaging can be found elsewhere (Nakatani et al., 2019). After averaging, the time intervals were a uniform 2min in duration. For each participant, timeseries were computed for NoTs and %CC. Event time-locked averaging was also applied to %MW and two EEG measures which we explain in the following sections.

The grand averages over participants are shown in Figure 2 of the Results section. The peaks in %MW at Bin0 reflect the segmentation criteria. Beyond Bin0, variation in %MW appears to be reduced over data bins. The result illustrates the averaging out of the non-time locked fluctuations in mind-wandering due to the segmentation procedure. This means that effects observed in the data will relate to the mind wandering in Bin0, rather than to some earlier or later mind-wandering events.

**Figure 2.**
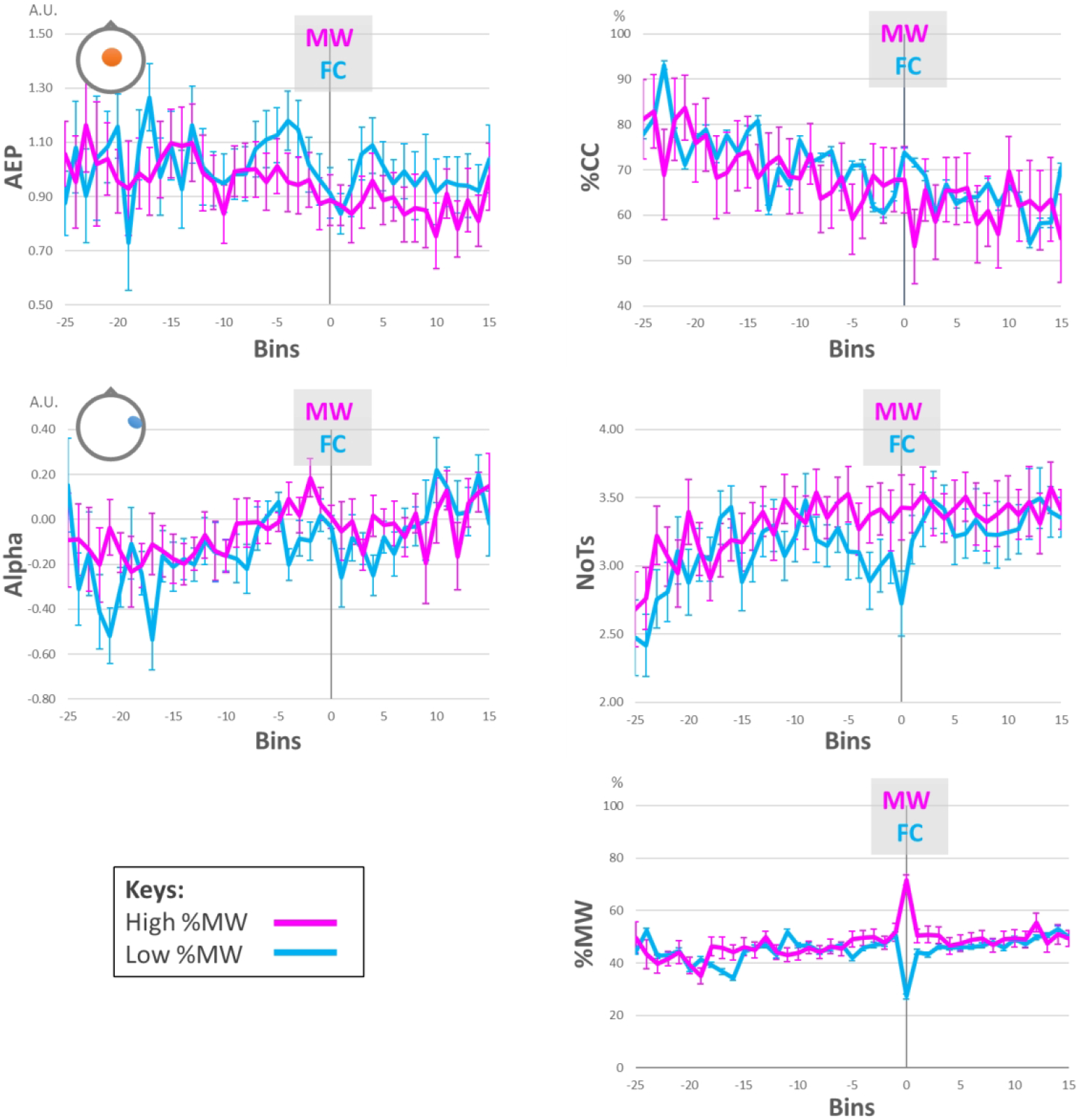
Event time-locked averages of AEP, Alpha, %CC, NoTs, and %MW. The horizontal axis represents time in data bins. Each bin interval equals 2 min. Bin0 starts from the point of subjective report of mind wandering (MW) in the high %MW condition (pink), and from that of task focus report (FC) in the low %MW condition (cyan). Error bars indicate SE. Head icons in the figures for AEP and Alpha indicate electrode location for the EEG measures.

### Preprocessing of EEG data

EEG data were pre-processed using Brain Vision Analyzer (version 2.1.2, Brain Products GmbH, Germany) and in-house Python scripts. Data were down-sampled from 2048 to 1024Hz, re-referenced to the average of all EEG channels, and filtered between 0.32 and 45.00Hz using a 2^nd^ order zero phase shift Butterworth filter. Independent component analysis (ICA, Infomax algorithm) was applied to the filtered data for artifact removal. EOG channels were included in the ICA analysis. Ocular, cardiac, and muscular artifact components were visually identified and excluded from the inverse ICA. After the cleaning, two channels (one for 2 participants) still showing poor signal quality were interpolated using adjacent channels.

### Computation of AEP amplitude

AEP has been reported as a complex of evoked components to auditory stimuli (Picton et al., 1974; Picton & Hillyard, 1974). Typically, the peak amplitude appears approx. 200-500ms from the onset of a pure tone stimulus, scalp distribution of the peak is central, and the peak frequency is situated in the theta-band (4- 8Hz). To test the timing, scalp distribution, and frequency of the AEP peak to the current stimulus (400Hz pure tone), we conducted time-frequency analysis using the EEG data of the odd-ball task, recorded from the same group of participants. The EEG data were segmented from the onset of standard (440Hz) tone, between −500 and 2000ms. Approx.100 segments were taken per participant. To obtain scalograms of instantaneous amplitude between 1 and 40Hz in logarithmic steps, the complex Morlet wavelet (no. of cycles = 5) was applied to each of the segments. The scalograms were normalized and averaged per participant. To the group data, t-tests were applied to test the significance of the amplitude against 0 at each data point in the scalogram. Type-I error was corrected using the false discovery rate (FDR) method (Benjamini & Hochberg, 1995). FDR was applied to the p-values of all data points over the 59 EEG electrodes. A more extended description of the procedure is available elsewhere (Bernhard, 2018). We found that amplitude in the frequency band between 4 and 8Hz, in time between 130 and 430ms from the stimulus onset, and in 7 central electrodes (3Z, 4Z, 5Z, 4L, 5L, 4R, 5R in the WaveGuard Duke64 system, cf. Appendix Fig. A2) showed p(FDR-corrected) < 0.05. The frequency, latency, and scalp distribution matched those in previous AEP studies (Picton et al., 1974; Picton & Hillyard, 1974), so these parameters were employed here to extract the AEP values used as GDP measures in the subsequent tone counting task.

EEG from the tone counting task was band-passed between 4 and 8Hz using in-house Python code for band-pass filtering, which combines a Butterworth filter (scipy.signal.butter, 12dB) and a zero-phase shift filter (scipy.signal.filtfilt). Instantaneous amplitude was computed using the complex-valued analytic signal from the Hilbert transform of the band-passed signal. The amplitude signal was segmented between −500 and 1000ms from a tone onset. Prior to averaging the segments, each segment was normalized using mean and standard deviation of the pre-stimulus period. Per trial, 20-24 segments were averaged. Based on the peak timing and scalp distribution parameters, amplitude between 130 and 430ms of the 7 central electrodes were chosen and averaged per trial per participant, and used as the AEP amplitude measure.

### Computation of alpha-band amplitude

EEG alpha-band activity (8-12Hz) is chosen as measure of on-going brain activity. Amplitude of the activity is known to be sensitive to differences in mental state, notably between resting and task states. (Chen et al., 2008; Compton et al., 2019; Gruberger et al., 2011; Marino et al., 2019). Accordingly, we may expect the amplitude to change over time as participants’ mental state fluctuates between task-focused and mind wandering states. We band passed the EEG between 8-12Hz using the above mentioned in-house band-pass filter for each available electrode (59ch). To the band-passed signals, Hilbert transform was applied to compute instantaneous amplitude. From the amplitude, we cut 2000ms periods, starting from the onsets of the 20-24 tone-stimuli within a trial. We averaged the amplitude of these periods at trial level to obtain timeseries of the single trial amplitudes.

Of the 59 signals, those with sufficient variability for the main analysis were selected. To avoid circular analysis, the channels were chosen based on the event-time locked average from an intermediate-level %MW report, 40≦%MW<60. The averaging was done in the same way as for %CC and NoTs. On average, 10.58 segments were averaged per participant. The procedure yielded average timeseries data (Appendix Figure A3) different from those used for the main analysis, which were averaged from high and low %MW (≧ 70 and < 30) reports, respectively (Figure 2).

The variability was tested at each electrode, fitting a mixed effect model, Alpha ∼ Time (in data bins, fixed effect factor) + participants (random effect factor) to the mid-level %MW segment, using the R package, lme4 (Bates, Mächler, Bolker, & Walker, 2014). The effect of Time was estimated using the chi-squared (χ^2^) statistic of ANOVA between the model and its null model: Alpha ∼ participants. The test was applied to all EEG electrodes. Type-I error over the univariate tests was corrected over the electrodes using the FDR method. Result showed a sizable effect of Time in two adjacent right temporal electrodes (3RB and 3RC; χ^2^(1) = 11.23, p(FDR-corrected) = 0.03, η^2^ = 0.43, power = 0.38 and χ^2^(1) = 12.33, p(FDR-corrected) = 0.02, η^2^ = 0.44, power = 0.34, respectively. The electrode locations are partially matching to those of tau activity (auditory resting alpha) studies, which involve bilateral temporal electrodes (Lehtelä et al., 1997; Tiihonen et al., 1991; Weisz et al., 2011). Given the electrodes are adjacent, the timeseries were averaged over the channels. From here on, this average is referred to simply as *Alpha*.

### Event time-locked averaging of AEP and Alpha

Event time-locked averaging was applied to the trial series of AEP and Alpha (Figure 1, middle). In each measure, the timeseries was segmented from the high- and from the low %MW reports, between −50 and +32min. The procedure of the averaging was same as that for NoTs and %CC. The averaging generated timeseries between Bins-25 to +15 from Bin0, for the high- and low %MW reports separately.

Grand average AEP and Alpha are shown in Figure 2 of the Results section. Temporally local features, peaks and troughs, are observable in the figure. However, analyzing such features in isolation may bias our understanding. Their selection is arbitrary and extrinsic to the dynamics of the timeseries. Therefore, rather than pre-selected temporal features, we included entire segments in the main data analysis.

### Models and main data analysis

We investigated whether the rate of thought generation and cognitive control are independently modulated by the amplitude of both AEP and Alpha. A set of models were built, starting from an autoregression model that predicts the current state of NoTs and %CC using their own past states. If past states of the neural variables improve the prediction of the current states beyond the autoregressive predictors, these neural variables are considered as stochastically causal predictors (Granger, 1969; Pearl, 2000). The following is step-by-step explanation of our approach, starting from the autoregression model. For NoTs, the autoregression model can be written as

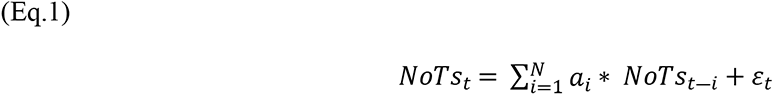

NoTs at a given time point *t*, *NoTs_t_*, is predicted by the weighted sum of own past states *NoTS_t-i_*, where *i* is the index for time regression and *a_i_* is the weight. The 𝜀_t_ is the error term. The past states were summed over *N*. To simplify the model, we temporarily set *N=1*. The simplified model is

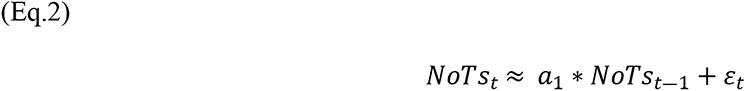

To the autoregression model, two neural predictors, *Alpha_t-1_* and *AEP_t-1_*, were added as follows.

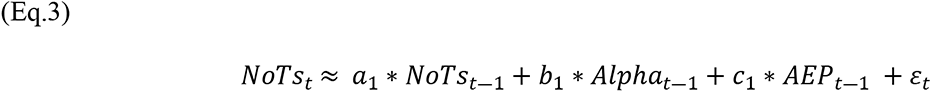

To the 3-predictor model, two more neural predictors, *Alpha_t_* and *AEP*_t_, were added. The predictors estimate neural effects without time difference from the response variable.

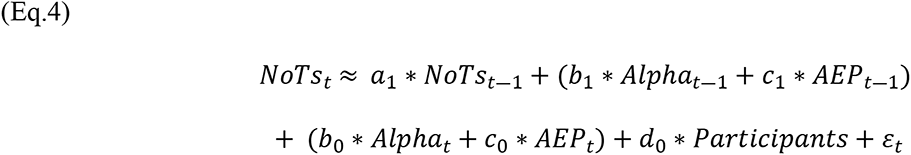

Parentheses are added for readability. A categorical variable *Participants* was added to account for individual differences. The same formulation was applied to predict %CC at time *t*, %*CC_t_*.

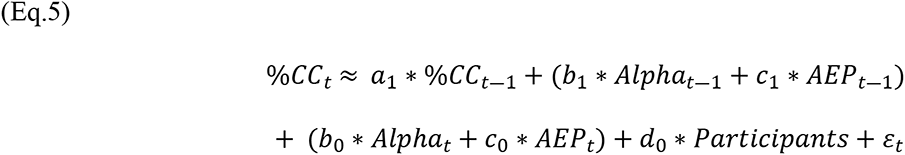

NoTs and %CC are two parallel data series in the tone counting task, which could covary. To account for the covariance, we chose a multivariate method to estimate the contribution of each predictor. The model for the estimation may be written as pseudo R code.

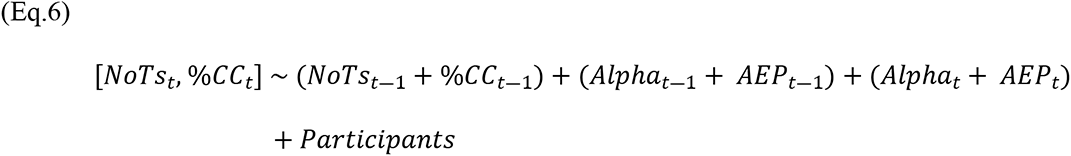

The multivariate linear regression method estimates the contribution of the autoregression, past-neural, present-neural predictors and the individual difference term to the prediction of the present state of NoTs and %CC, taking into account possible correlations among the variables. Reduced models can be obtained from this model (Eq.6) to evaluate the stochastic causality of Alpha and AEP effects; a model without *Alpha*_𝑡―1_ and another without *AEP_t-1_*. If the reduced models predict the response variables worse than the full model did, the contribution of the omitted neural predictor is not negligible. I.e., the comparison gives the basis for attributing stochastic causality to the effect.

The above models set the time regression as *t-1*, which means the most recent past state represents the effect of all past states. This would be appropriate if we could assume that the most recent past state has always the strongest effect. However, as discussed in the Introduction, we assumed that the effect timing could vary depend on the task-focused and mind-wandering states. Thus, the order of regression was parametrized using indices *i, j, k,* and *l*.

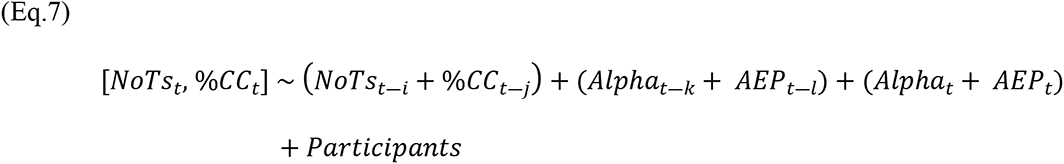

The parameters vary systematically between 1 and 10 (given the data bin size of 2min, this corresponds to times between 2 and 20min. The range was decided by the length of the data).

The full combination of the 4 parameters yields a number of 10^4^ models to fit. The number of models was brought down in two steps. In the first step, *i* and *j* were searched between 1 and 10 for the highest autoregressive effects, while *k* and *l* were fixed to *k=1* and *l=1*. The larger autoregression terms were desirable since they make the criterion for stochastic causality more strict. The resulting 10 by 10 = 100 models were fit to the high- and low %MW segments, separately. Upon fitting, a data window was applied; the window for response variables, *NoTs_t_*and %𝐶𝐶_t_, was ±15 bins from Bin0 (which starts from *t*, the moment of the %MW report) based on the results of subjective report and task measure in our previous study (Nakatani et al., 2019). The same data window was applied to the present neural predictors. For the autoregression and past neural state predictors, the size of the data window was the same, however, the window was shifted bin-by-bin, up to 10bins (20min) to the past, as indicated by *i* and *j.* Of the 100 combinations of *i* and *j*, those with the highest autoregressive effects were selected using adjusted-R^2^. In case of tie, the smallest *i* an*d j* were chosen e.g., if (*i=1, j=*2) and (*i=2, j=*2) gave the highest adjusted-R^2^, (*i=1, j=*2).

In the second step, *k* and *l* were systematically varied between 1 and 10, while the autoregression terms were fixed with *i=i\** and *j=j\**, where *i\** and *j\** were those selected in the first step. In this manner, the number of full models were reduced from 10^4^ to 10^2^. The full model with *i\** and *j\** in pseudo R code is

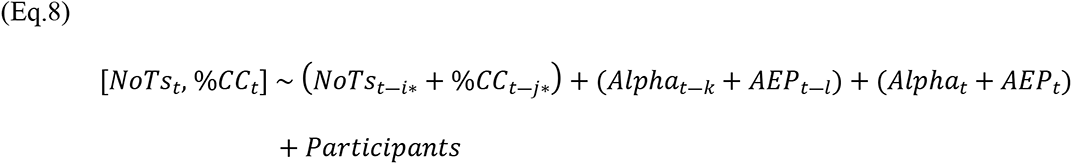

The models were fit to the high- and low %MW segment separately. To test the stochastic causality, 2 reduced models, one without the past Alpha and the other without the past AEP term, were also fit to the data. The reduced models are:

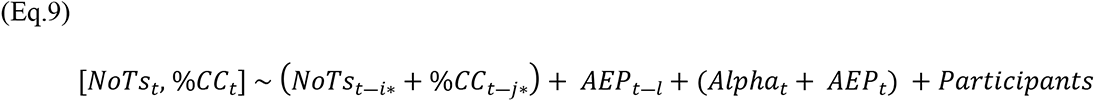

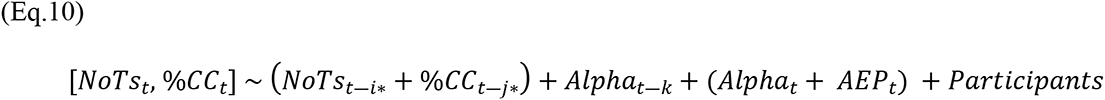

The Pillai test was used to estimate statistical significance of the difference between the fits of the full and reduced models. The 100 comparisons for 2 reduced models yielded 200 tests. The R implementation of Pillai test provides the estimated F-statistic with corresponding probability. We corrected for Type-I errors over the 200 tests by using the FDR method. An arbitrary threshold (p<0.01) was applied to the corrected probability, as criterion for the attribution of stochastic causality to the neural effects. The indices *k* and *l* reveal the timing of the effects, e.g., a stochastically causal effect of *Alpha*_t-𝑘_ at *k=4* on *NoTs_t_*suggests that (given the 2min interval of data bin) the neural activity between 6-8min *before* the report of the high (or low) %MW had stochastically causal effect on the number of spontaneous thoughts 6-8 min later.

## Results

### Visual inspection of event time-locked average data

Event time-locked averages of the five measures AEP, Alpha, NoTs, %CC, and %MW, are available for visual inspection in Figure 2. Most show linear trends and local patterns, while %MW was essentially flat, except for the peaks at Bin0, which are, according to their condition, focused: %MW < 30; mind-wandering: %MW ≧ 70. Elsewhere %MW appears homogeneously distributed. Averaging has reduced non time-locked effects of %MW in the same way as averaging increases the signal-to-noise ratio in ERP. Thus, peaks and troughs in the other variables, AEP, Alpha, NoTs, and %CC, can be analyzed as time-locked to the mind-wandering report at Bin0.

AEP amplitude was higher in low- than in high %MW conditions most of the time. The difference, however, was reduced around the moment of subjective report, in Bin0. Alpha amplitude was higher in the high- than in low %MW conditions, mostly before the report. The %CC decreased with time. The task performance measure was high around the subjective report of task-focus (%MW <30 in Bin0), while it was low around the report of mind wandering (%MW ≧70 in Bin0). In contrast, while NoTs increased with time overall, there was a dip around the period task focus was reported. For all measures, preliminary analyses indicated a sizable conditions-by-time interaction and some of the bin-wise differences were reliable. However, they would be hard to interpret in isolation. An analysis based on such local differences and features relies only on part of the variance in the data and contributes little to our understanding of the dynamics. Instead, we proceed to the main analysis, which takes all of the variance in the data segments into account.

Prior to the model fit, multi-variate outliers were removed using Mahalanobis distance among NoTs, %CC, Alpha and AEP data; datapoints with Mahalanobis distance > 20 were excluded from the model fit. Linear trends were removed to make the data quasi-stationary. This is to avoid bias in model estimation due to the trend. The result of the linear trend was reported in the section, *Notes on linear trends*.

The detrended high- and low %MW data segments were used in the step-wise model fit.

### Results of the Main Analysis

#### High %MW segments

The first step of model fitting indicated *i = 2* and *j = 5* for the highest autoregression effect (See *Notes on autoregressive effects*). For the second step, *i\** and *j\** were set as 2 and 5, respectively, and 100 models were generated over the combinations of *k* and *l*. The full- and corresponding reduced models were fit to the high %MW segments data. The difference between the full- and reduced models was tested using the Pillai test. Type-I error was corrected over the 200 tests using FDR method. Table1 shows the corrected probability of the estimated-F statistic of the Pillai score. With an arbitrary threshold of p<0.01, stochastic causality was suggested for past neural predictor *Alpha*_t-𝑘_ at *k =* 4 and 5 (Table 1a), while no causality was suggested for *AEP*_t-𝑙_ (Table 1b). Timing of the effect at *k =* 4 and 5 correspond to 6-8 and 8-10min before the high %MW report, respectively.

Table 2 list the estimated coefficients for the full model with *k=4 and 5*. (For both cases, the index for *AEP*_t-𝑙_ was *l=1.* With the choice, no crucial information was missed since the effect of *AEP*_t-𝑙_changed little over *l.*) The coefficient for the past neural effect *Alpha*_t-4_which was positive as a predictor for %𝐶𝐶_t_; coefficient = 8.16, the 95% Confidence Interval [2.5%, 97.5%] = [4.22, 12.11], t-value = 4.07. The effect was sizable compared with that of the autoregression term, %𝐶𝐶_t-5_; coefficient = 0.21, 95%CI = [0.12, 0.31], t-value = 4.33. By contrast, the effects of the present neural predictors *Alpha*_t_ and *AEP*_t_ to %𝐶𝐶_t_were negligible. For the prediction of *NoTs_t_*, no neural effect was noteworthy, while the autoregression effect was considerable; coefficient of *NoT*𝑠_t-2_= 0.24, 95%CI = [0.15, 0.33], t = 5.29. Results of *k=5* (Table 2b) showed the same pattern as *k=4*; The effect of *Alpha*_t-5_to %𝐶𝐶_t_was positive, and also the auto regression effects were positive.

**Table 1:**
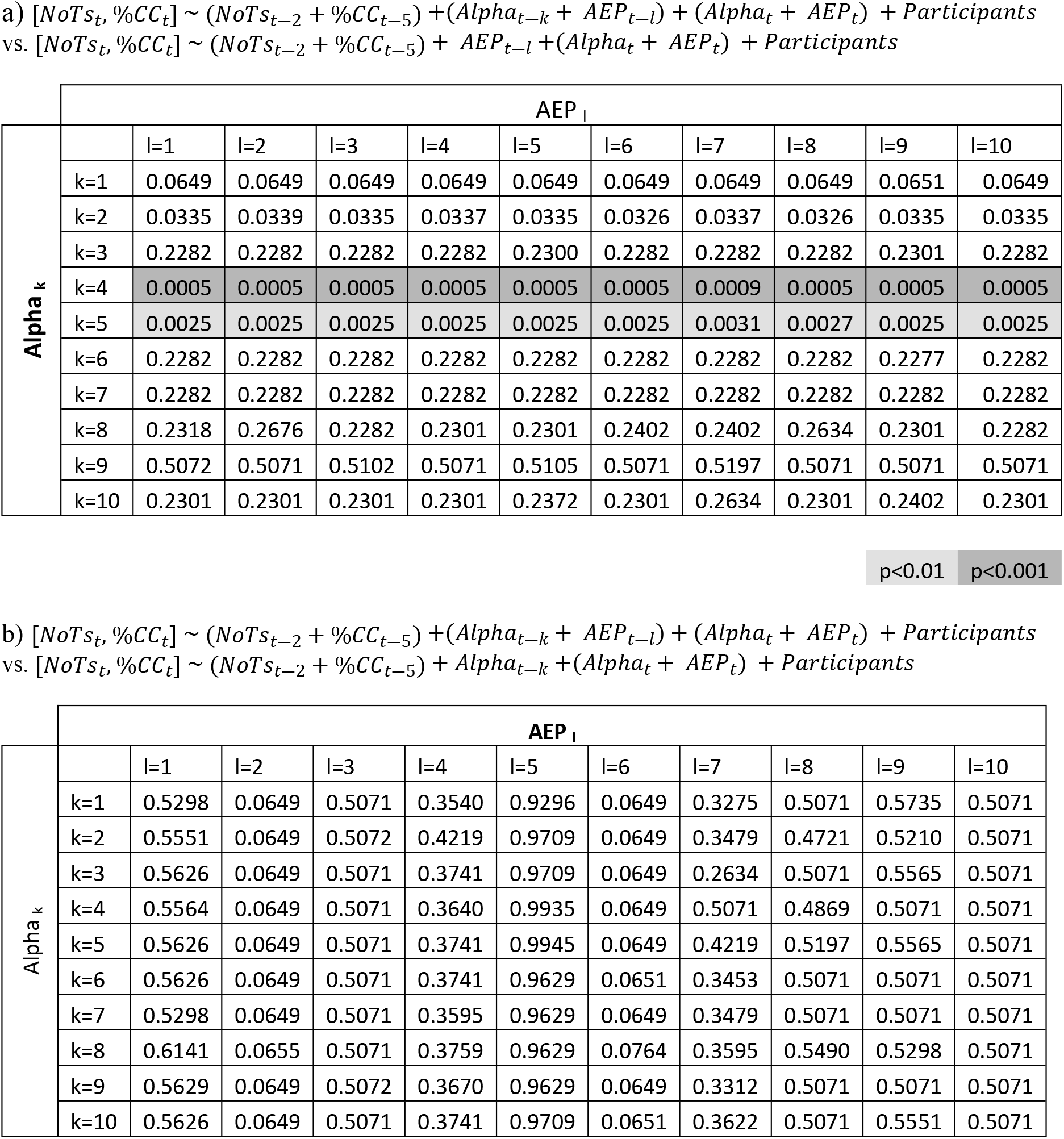
Effect of past neural states in High %MW condition. a) Model comparison tests the stochastic causality of past neural Alpha states, b) tests the stochastic causality of past neural AEP states. Pillai test was applied to test the difference between full and reduced models. Probability of estimated Fstatistic, DF = (1, 462) of Pillai score was shown after FDR correction.

**Table 2:** Estimated coefficients of the full model for High %MW segments, separately for *Alpha_t-4_* (a) and *Alpha_t-5_* (b).

a) $[NoTs_t, \%CC_t] \sim (NoTs_{t-2} + \%CC_{t-5}) + (Alpha_{t-4} + AEP_{t-1}) + (Alpha_t + AEP_t) + Participants$
|  | Predictors | Coefficients | Confidence interval |  | t-value | Pr(>\|t\|) |
| --- | --- | --- | --- | --- | --- | --- |
|  |  |  | 2.50% | 97.50% |  |  |
| Response var. = %CC<sub>t</sub> | Intercept | -4.39 | -7.06 | -1.72 | -3.23 | 0.001 |
|  | NoTs<sub>t-2</sub> | -0.37 | -3.81 | 3.07 | -0.21 | 0.834 |
|  | %CC<sub>t-5</sub> | 0.21 | 0.12 | 0.31 | 4.33 | 0.000 |
|  | Alpha<sub>t-4</sub> | 8.17 | 4.22 | 12.11 | 4.07 | 0.000 |
|  | AEP<sub>t-1</sub> | 0.93 | -5.01 | 6.87 | 0.31 | 0.760 |
|  | Alpha<sub>t</sub> | -0.44 | -4.16 | 3.28 | -0.23 | 0.815 |
|  | AEP<sub>t</sub> | 4.78 | -1.07 | 10.64 | 1.61 | 0.109 |
|  | Participants | 0.39 | 0.17 | 0.61 | 3.46 | 0.001 |
| Response var. = NoTs<sub>t</sub> | Intercept | -0.02 | -0.09 | 0.05 | -0.48 | 0.631 |
|  | NoTs<sub>t-2</sub> | 0.24 | 0.15 | 0.33 | 5.29 | 0.000 |
|  | %CC<sub>t-5</sub> | 0.00 | 0.00 | 0.00 | 0.54 | 0.588 |
|  | Alpha<sub>t-4</sub> | 0.09 | -0.01 | 0.19 | 1.71 | 0.088 |
|  | AEP<sub>t-1</sub> | -0.09 | -0.24 | 0.06 | -1.17 | 0.244 |
|  | Alpha<sub>t</sub> | 0.07 | -0.03 | 0.16 | 1.39 | 0.164 |
|  | AEP<sub>t</sub> | 0.11 | -0.04 | 0.27 | 1.49 | 0.137 |
|  | Participants | 0.00 | 0.00 | 0.01 | 1.38 | 0.169 |

b) $[NoTs_t, \%CC_t] \sim (NoTs_{t-2} + \%CC_{t-5}) + (Alpha_{t-5} + AEP_{t-1}) + (Alpha_t + AEP_t) + Participants$
|  | Predictors | Coefficients | Confidence interval |  | t-value | Pr(>\|t\|) |
| --- | --- | --- | --- | --- | --- | --- |
|  |  |  | 2.50% | 97.50% |  |  |
| Response var. = %CC<sub>t</sub> | Intercept | -4.48 | -7.16 | -1.80 | -3.28 | 0.001 |
|  | NoTs<sub>t-2</sub> | -0.60 | -4.06 | 2.86 | -0.34 | 0.732 |
|  | %CC<sub>t-5</sub> | 0.20 | 0.11 | 0.30 | 4.06 | 0.000 |
|  | Alpha<sub>t-5</sub> | 6.92 | 3.00 | 10.85 | 3.46 | 0.001 |
|  | AEP<sub>t-1</sub> | 2.35 | -3.63 | 8.32 | 0.77 | 0.440 |
|  | Alpha<sub>t</sub> | 0.46 | -3.22 | 4.14 | 0.25 | 0.807 |
|  | AEP<sub>t</sub> | 3.48 | -2.40 | 9.37 | 1.16 | 0.246 |
|  | Participants | 0.40 | 0.18 | 0.62 | 3.55 | 0.000 |
| Response var. = NoTs<sub>t</sub> | Intercept | -0.02 | -0.09 | 0.05 | -0.50 | 0.619 |
|  | NoTs<sub>t-2</sub> | 0.24 | 0.15 | 0.33 | 5.23 | 0.000 |
|  | %CC<sub>t-5</sub> | 0.00 | 0.00 | 0.00 | 0.40 | 0.688 |
|  | Alpha<sub>t-5</sub> | 0.09 | -0.01 | 0.19 | 1.76 | 0.079 |
|  | AEP<sub>t-1</sub> | -0.07 | -0.23 | 0.08 | -0.95 | 0.343 |
|  | Alpha<sub>t</sub> | 0.08 | -0.02 | 0.17 | 1.59 | 0.113 |
|  | AEP<sub>t</sub> | 0.10 | -0.05 | 0.25 | 1.29 | 0.198 |
|  | Participants | 0.00 | 0.00 | 0.01 | 1.41 | 0.159 |

The above results show that Alpha activity 6-10min before the high %MW report had a stochastically causal and task-*positive* effect on the %CC. The rest of neural effects did not show notable contribution to predict the %CC and NoTs.

#### Low %MW segments

Results of the first-stage model fit suggested *i= 1* and *j= 9* for the highest autoregression effect. Thus, *i\** and *j\** of the model in the second stage was set as 1 and 9, respectively. With the selected autoregression terms, 100 models were generated over the combination of *k* and *l* between 1 and 10. The full- and reduced models were fit to the data in the low %MW segment. The Pillai test was applied in the same way for the analysis of high %MW segment data, to assess the stochastic causality of the neural effects. As shown in Table 3, the test results suggested the causality for *Alpha*_t-𝑘_ at *k =* 3, while no causality was suggested for *AEP*_t-𝑙_ . The corresponding timing for *Alpha*_t-3_ was 4-6min before the low %MW report.

**Table 3:** Effect of past neural states in Low %MW condition. a) Model comparison tests the stochastic causality of past neural Alpha states, b) tests the stochastic causality of past neural AEP states. Pillai test was applied to test the difference between full and reduced models. Probability of estimated Fstatistic, DF = (1, 395) of Pillai score was shown after FDR correction.

|  |  | AEP<sub>l</sub> |  |  |  |  |  |  |  |  |  |
| --- | --- | --- | --- | --- | --- | --- | --- | --- | --- | --- | --- |
| Alpha<sub>k</sub> |  | l=1 | l=2 | l=3 | l=4 | l=5 | l=6 | l=7 | l=8 | l=9 | l=10 |
|  | k=1 | 0.9552 | 0.9552 | 0.9552 | 0.9629 | 0.9552 | 0.9552 | 0.9552 | 0.9552 | 0.9935 | 0.9552 |
|  | k=2 | 0.9935 | 0.9935 | 0.9935 | 0.9935 | 0.9935 | 0.9935 | 0.9935 | 0.9935 | 0.9935 | 0.9935 |
|  | k=3 | 0.0014 | 0.0014 | 0.0014 | 0.0014 | 0.0014 | 0.0014 | 0.0014 | 0.0014 | 0.0014 | 0.0014 |
|  | k=4 | 0.9935 | 0.9935 | 0.9935 | 0.9935 | 0.9935 | 0.9935 | 0.9935 | 0.9935 | 0.9935 | 0.9935 |
|  | k=5 | 0.0952 | 0.0952 | 0.0988 | 0.0952 | 0.0952 | 0.0952 | 0.0952 | 0.0952 | 0.0952 | 0.0952 |
|  | k=6 | 0.3069 | 0.3222 | 0.3222 | 0.2485 | 0.3038 | 0.3069 | 0.3069 | 0.3222 | 0.2898 | 0.3069 |
|  | k=7 | 0.5537 | 0.5579 | 0.5719 | 0.6140 | 0.5683 | 0.5579 | 0.5579 | 0.5286 | 0.5579 | 0.5579 |
|  | k=8 | 0.9935 | 0.9935 | 0.9935 | 0.9935 | 0.9935 | 0.9935 | 0.9935 | 0.9935 | 0.9935 | 0.9935 |
|  | k=9 | 0.0636 | 0.0453 | 0.0636 | 0.0636 | 0.0636 | 0.0636 | 0.0636 | 0.0928 | 0.0636 | 0.0636 |
|  | k=10 | 0.7600 | 0.7526 | 0.8904 | 0.7585 | 0.7585 | 0.7492 | 0.7585 | 0.7600 | 0.7492 | 0.7585 |

| AEP<sub>l</sub> |  |  |  |  |  |  |  |  |  |  |  |
| --- | --- | --- | --- | --- | --- | --- | --- | --- | --- | --- | --- |
| Alpha<sub>p</sub> |  | l=1 | l=2 | l=3 | l=4 | l=5 | l=6 | l=7 | l=8 | l=9 | l=10 |
|  | k=1 | 0.9935 | 0.7492 | 0.6698 | 0.7492 | 0.9935 | 0.7585 | 0.9935 | 0.4371 | 0.1102 | 0.9935 |
|  | k=2 | 0.9935 | 0.7492 | 0.6698 | 0.7492 | 0.9935 | 0.7585 | 0.9935 | 0.4371 | 0.0952 | 0.9935 |
|  | k=3 | 0.9935 | 0.7492 | 0.6698 | 0.7492 | 0.9935 | 0.7600 | 0.9935 | 0.4435 | 0.1077 | 0.9935 |
|  | k=4 | 0.9935 | 0.7492 | 0.6698 | 0.7492 | 0.9935 | 0.7585 | 0.9935 | 0.4371 | 0.0952 | 0.9935 |
|  | k=5 | 0.9935 | 0.7492 | 0.7492 | 0.7492 | 0.9935 | 0.7803 | 0.9935 | 0.4535 | 0.0952 | 0.9935 |
|  | k=6 | 0.9935 | 0.7585 | 0.7296 | 0.6835 | 0.9935 | 0.7585 | 0.9935 | 0.4562 | 0.0952 | 0.9935 |
|  | k=7 | 0.9935 | 0.7492 | 0.6914 | 0.7585 | 0.9935 | 0.7585 | 0.9935 | 0.4099 | 0.0952 | 0.9935 |
|  | k=8 | 0.9935 | 0.7492 | 0.6698 | 0.7492 | 0.9935 | 0.7585 | 0.9935 | 0.4371 | 0.0952 | 0.9935 |
|  | k=9 | 0.9935 | 0.5083 | 0.6698 | 0.7526 | 0.9552 | 0.7492 | 0.9935 | 0.5683 | 0.0952 | 0.9935 |
|  | k=10 | 0.9935 | 0.7492 | 0.7492 | 0.7492 | 0.9935 | 0.7492 | 0.9935 | 0.4392 | 0.0952 | 0.9935 |

Table 4 lists the estimated coefficients for the predictors at *k =* 3. Similar to Table 2, index *l* for *AEP*_t-𝑙_ was set to *1* to simplify the presentation of the results. The coefficient for the past neural effect term *Alpha*_t-3_for %𝐶𝐶_t_was large; coefficient = −10.04, 95% CI = [−14.6, −5.43], t-value = -4.29, relative to that of autoregression term, %𝐶𝐶_―9_, coefficient = 0.15, 95%CI = [0.07, 0.24], t-value = 3.51. Thus, the past neural effect could be qualified for stochastic causality similar to that in the high %MW segment, although the sign was *negative*. Moreover, in the low %MW segments, the effect of present Alpha was noticeable; coefficient of *Alpha*_t_= 7.38, 95% CI = [2.77, 12.00], t-value = 3.15.

**Table 4:** Estimated coefficients of the full model for Low %MW condition for *Alpha_t-3_*. [*NoTs_t_* %*CC_t_*] ~ (*NoTs_t-1_* + %*CC_t-9_*) +(*Alpha_t-3_* + *AEP_t-1_*) + (*Alpha_t_* + *AEP_t_*) + *Participants*

|  | Predictors | Coefficients | Confidence interval |  | t-value | Pr(>\|t\|) |
| --- | --- | --- | --- | --- | --- | --- |
|  |  |  | 2.50% | 97.50% |  |  |
| Response var. = %CC<sub>t</sub> | Intercept | -6.54 | -10.27 | -2.82 | -3.45 | 0.0006 |
|  | NoTs<sub>t-1</sub> | 4.83 | 0.81 | 8.86 | 2.36 | 0.0188 |
|  | %CC<sub>t-9</sub> | 0.15 | 0.07 | 0.24 | 3.51 | 0.0005 |
|  | Alpha<sub>t-3</sub> | -10.04 | -14.64 | -5.43 | -4.29 | 0.0000 |
|  | AEP<sub>t-1</sub> | 1.85 | -4.59 | 8.30 | 0.57 | 0.5721 |
|  | Alpha<sub>t</sub> | 7.38 | 2.77 | 12.00 | 3.15 | 0.0018 |
|  | AEP<sub>t</sub> | -1.01 | -7.61 | 5.58 | -0.30 | 0.7628 |
|  | Participants | 0.57 | 0.26 | 0.88 | 3.65 | 0.0003 |
| Response var. = NoTs<sub>t</sub> | (Intercept) | -0.07 | -0.15 | 0.02 | -1.50 | 0.1343 |
|  | NoTs<sub>t-1</sub> | 0.12 | 0.03 | 0.21 | 2.57 | 0.0107 |
|  | %CC<sub>t-9</sub> | 0.00 | 0.00 | 0.00 | -0.27 | 0.7887 |
|  | Alpha<sub>t-3</sub> | 0.10 | 0.00 | 0.21 | 1.89 | 0.0592 |
|  | AEP<sub>t-1</sub> | 0.03 | -0.12 | 0.18 | 0.37 | 0.7105 |
|  | Alpha<sub>t</sub> | -0.17 | -0.28 | -0.07 | -3.22 | 0.0014 |
|  | AEP<sub>t</sub> | 0.04 | -0.11 | 0.19 | 0.51 | 0.6115 |
|  | Participants | 0.01 | 0.00 | 0.01 | 2.06 | 0.0401 |

In contrast, *Alpha*_t-3_ predicted little of *NoTs_t_*; coefficient = 0.10, 95%CI = [0.00, 0.21], t-value = 1.89, relative to that of the autoregressive term *NoT*𝑠_―1_ coefficient = 0.12, 95%CI = [0.03, 0.21], t-value = 2.57. Besides, the effect of present Alpha was noticeable; coefficient of *Alpha*_t_ = −0.17, 95% CI = [−0.28, −0.07], t-value = -3.22.

The above results show that Alpha activity 4-6min before the low %MW report had a stochastically causal and task-*negative* effect on %CC. Moreover, the results show that the Alpha activity correlated without delay (in the current time scale), positively with %CC, while negatively with NoTs in the low %MW segment.

#### Notes on autoregressive effects

The indices for autoregression terms, *i\** and *j\**, were selected to make the criterion for stochastic causality as strict as possible, and to reduce the number of full models. We chose *i* with the highest adjusted-R^2^ for NoTs, and *j* with the highest adjusted-R^2^ for %CC. The selected *i\** and *j\** were 2 and 5 for the high-, and 1 and 9 for the low %MW segments. The timing of the effects, in particular those for the autoregression effect of %CC, were further in the past than might have been expected. As a reality check, we also fit full models with autoregression terms with *i*= 1 and *j*=1 to the data. The results showed that the contribution of the terms were slightly less than those with *i\** and *j\**; e.g., %𝐶𝐶_t-1_had coefficient= 0.11, 95%CI=[0.01, 0.21], t=2.17 (prob.=0.031), in comparison with %𝐶𝐶_t-9_ coefficient = 0.15, 95%CI=[0.07, 0.24], t=3.51 (prob.=0.0005) for the low %MW segment (See Appendix 4 for more results). The observations suggest that the *i\** and *j\** gave a stricter baseline for causality. Thus, we conclude the *i\** and *j\** chosen are reasonable for the purpose in the current study.

#### Notes on linear trend

Prior to the model fit, the linear trend was removed to make the data segment quasi-stationary. However, the linear trend is useful to check if the %CC and NoTs were negatively correlated during the task. We expected the %CC to decrease, while NoTs to increase with time, thus, a negative and positive linear trend for %CC and NoTs, respectively.

The linear trend was estimated applying a mixed-effect model, Y ∼ Time (fixed variable) + participants (random variable) to %𝐶𝐶_t_and *NoTs_t_*in of the low- and high %MW segment, separately. In the low %MW segment, the linear trend was negative for %CC; Linear coef.= -23.27, the 95%CI= [−32.20, −14.34], t= -4.97, and positive for NoTs; Linear coef.= 0.55, the 95%CI= [0.31, 0.78], t= 4.42. In the high %MW segment, the linear coefficient for %𝐶𝐶_t_was again negative; Linear coef.= −15.79, the 95%CI= [−23.66, -7.84], t= -3.80. For *NoTs_t_*, the positive trend was not strong; Linear coef.= 0.18, the 95%CI= [− 0.03, 0.39], t= 1.66. Overall, the results suggest that the %CC and NoTs negatively correlated during the task. The weak linear trend in the high %MW segment might suggests that the NoTs were near their ceiling; the number of thoughts varied around 3.40 which is near 4, the limit in working memory capacity according to Luck and Vogel (1997). Moreover, this is above the 90% level of the standardized number of thoughts observed in our previous study (Nakatani et al., 2019). In contrast, the number varied around 2.72 in the low %MW segment.

Moreover, we checked the linear trend of *AEP*_t_ and *Alpha*_t_. In the low %MW segment, the AEP amplitude decreased with time; Linear coef.= −0.28, the 95%CI= [−0.43, −0.14], t= -3.75, while the Alpha amplitude increased; Linear coef.= 0.42, the 95%CI= [0.21 0.64], t= 3.75. Similarly, in the high %MW segment, AEP amplitude decreased; Linear coef.= −0.37, the 95%CI= [−0.50, −0.24], t= −5.53, while Alpha amplitude increased with time; Linear coef.= 0.26, the 95%CI= [0.07, 0.46], t= 2.59.

The above results are in agreement with a common dynamic overall, which seesaws NoTs and %CC, and also Alpha and AEP. However the effect of *Alpha*_t-𝑘_(*k=*3, 4, and 5) on %CC shows that the seesaw was at times disrupted by transient neural activity related to cognitive control.

## Discussion

In the context of a tedious task, episodes of mind wandering are likely to occur. In the transition to mind wandering, the rate of thought generation and the availability of cognitive control for the task may both play a critical role. We hypothesized that both vary autonomously over time. These fluctuations eventually are manifested in subjective reports of mind wandering and related mental states (Christoff et al., 2016; Jaarsveld & Van Leeuwen, 2005) and in the complex patterns of transitions between these states (Nakatani et al., 2019). We investigated the dynamic properties of thought generation and cognitive control for task-focused and mind wandering states. Participants performed a tedious tone counting task for a 1.5 hour period. During this period, subjective reports of numbers of thoughts (NoTs) and percentage mind wandering (%MW) were recorded, along with the performance on the tone counting task (%CC) and two EEG signals, AEP and Alpha amplitude.

In our experiment, NoTs increased and %CC decreased steadily over time during the task. These opposing linear trends might suggest that thought generation and cognitive control are yoked in a common dynamic that seesaws the rate of thought generation and the availability of cognitive control. However, such a seesaw dynamic would only explain part of the complex fluctuations in NoTs and %CC. In the data segment towards mind wandering, the autoregressive effect on NoTs was strongest at lag *i\** = 2, which corresponds to 2-4min before a report of mind wandering (according to our criterion of reported %MW ≧ 70), and the effect on %CC was strongest at lag *j\** = 5, corresponding to 8-10min before the mind wandering report. Similarly, autoregressive effects on NoTs and %CC peaked at 0-2min and 16-18min, respectively, before a task focus report (%MW < 30). These results confirm our reasoning that thought generation and cognitive control have distinctive contributions to the dynamics of mind wandering during the execution of a boring task.

It may be counterintuitive that the timing of the highest autoregressive effect was not at lag=1, except for NoTs towards task-focused reports. It means that the most recent past event was not the strongest predictor for the current state. The results show that the crucial autoregressive effects are spread over minutes. There is a hint of periodicity in the data. However, we do not have a clear interpretation for the precise distribution of the lagged effects.

We next analyzed how the two EEG measures, AEP and Alpha amplitude, influenced the fluctuations in NoTs and %CC. AEP showed no notable effect on any of our measures. It is generally considered that AEP amplitude is sensitive to the level of attention for information processing of incoming auditory stimuli (Picton et al., 1974; Picton & Hillyard, 1974). However, the level of attention required to process the tone stimuli themselves may not be very high. Thus, fluctuations of AEP amplitude might have not been a critical factor on the performance of the tone counting task. Alternatively, the sensitivity of the evoked potentials (EPs) to sustained attention may have been low. In a study of sustained attention using visual stimuli, visual evoked potential (VEP) amplitude was less sensitive to sustained attention than visual alpha-band amplitude. (Dockree et al., 2007). In the current dataset, however, the sensitivity between the AEP amplitude (full band) and the auditory alpha-band power did not differ (Bernhard, 2018). Thus, the level of attention required by the tone counting task is the more likely explanation for the lack of an AEP effect.

In contrast with AEP, a reliable effect of Alpha on %CC was obtained. The Alpha effect appeared 6-10min before a report of mind wandering, whereas it appeared 4-6min before a report of task focus. Interestingly, the direction of the effect was positive for the mind-wandering and negative for the task-focused reports. Both positive and negative effects of alpha-band amplitude have been reported in previous studies of resting-state alpha activity (Kam et al., 2022). A switch of effect direction within a task was reported in BOLD-fMRI signal during an autobiographical memory task (Andrews-Hanna et al., 2014). This study focused on the activity of the default-mode network (DMN), which generally shows higher activity in rest- than in task conditions, i.e., task negativity (Greicius et al., 2003; Raichle, 2015). However, during an autobiographical memory task, subregions of the DMN, i.e., dorsomedial prefrontal cortex, temporal-parietal junction, lateral temporal cortex, and temporal pole showed higher BOLD levels than in resting conditions. This effect was further increased when the task required detailed examination of autobiographic memory (Andrews-Hanna et al., 2014). The authors argued that the switch from task-negative to positive in the DMN regions reflects a shift in mental focus from external events to autobiographical memory. In a review, Buckner and Dinicola (2019) linked task positivity of the DMN subregions to “internally generated information.” In terms of our study, this implies that as the mind begins to wander, the mental focus shifts from the external auditory stimuli of the task to internally generated thoughts. While the directional changes in the fMRI and EEG studies may not be readily comparable, both studies at least suggest that neural effects become more variable as the mental process incorporates “internally generated information.”

Neither Alpha nor AEP had a significant effect on NoTs. It is surprising that the ongoing alpha activity did not predict NoTs, particularly as EEG alpha band activity has been associated with resting state and default mode network activity (Chen et al., 2008; Compton et al., 2019; Gruberger et al., 2011). Previous studies have focused on alpha activity from posterior electrodes. However, in our data the alpha band activity from parietal, occipital, and left temporal regions did not show sufficient variance to qualify as model predictors for NoTs. We were therefore unable to identify any neural predictors of fluctuations of thought production on the timescale of minutes.

The current EEG results have potential applications for mind wandering in real-life. EEG recording using a mobile system with a small number of channels, e.g., one channel in the right temporal scalp location, may be sufficient to obtain the necessary signals. The current results specified the timing of possible EEG marker signal, 4-10 minutes before the subjective report of mind wandering. The signal could be combined with bandpass filtering, which can be performed (quasi) in real-time, to develop a reliable marker for the prediction of mind wandering in various applied settings.

To envisage the wider implications of our findings, consider a typical experiment contrasting focused and mind wandering states. Suppose the study seeks, and finds, a unique neural correlate of mind wandering, which might even be used to predict transitions to mind wandering. Yet, without giving consideration to the possible independent contributions of thought production and cognitive control, this approach would fail to adequately characterize the transitions between focused and mind wandering states. On the map of mental states relevant for mind wandering (Christoff et al., 2016), the transitions would occur on the antidiagonal (Figure 3). In our view, this would be an oversimplification. The wider context of spontaneous mental process, in which mind wandering is situated, would be lost by correlating neural data directly with subjective reports of mind wandering.

**Figure 3.**
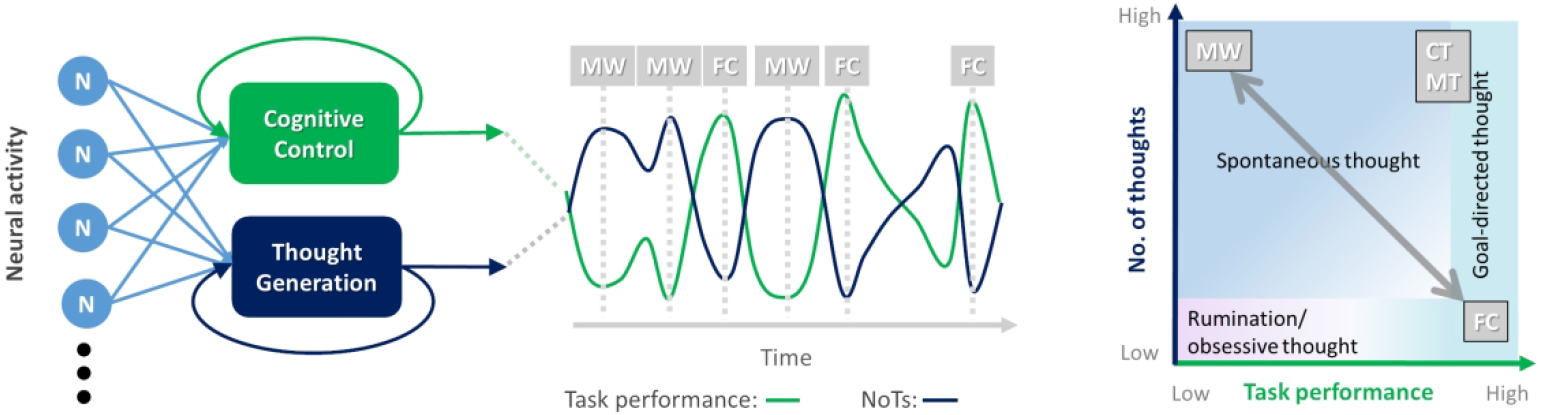
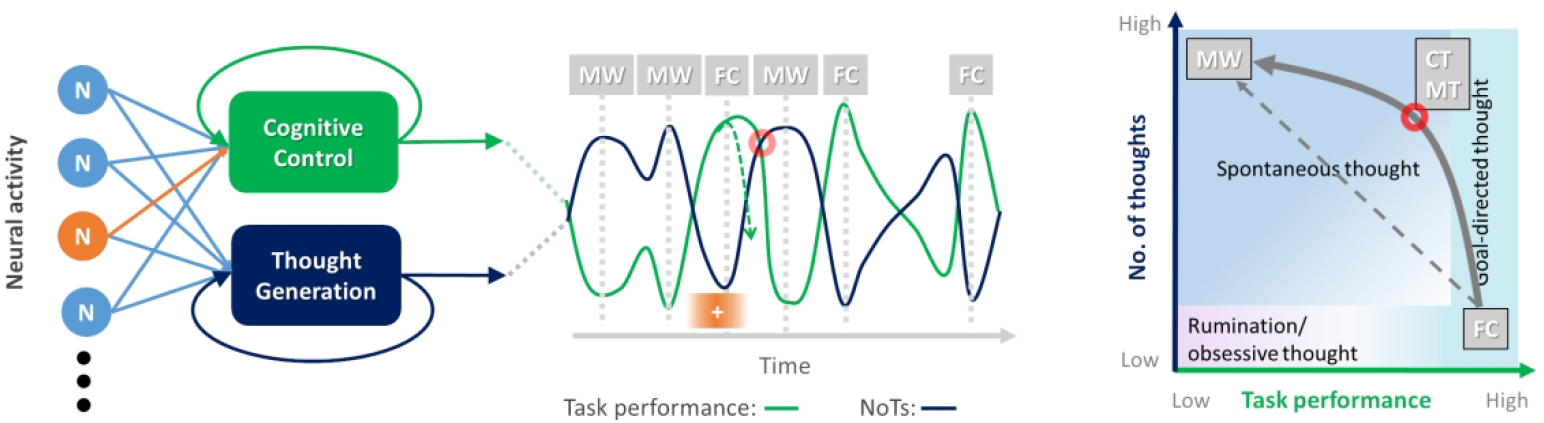
Illustration of possible dynamic neural effects on the path toward mind wandering. a) Neural activity sources are seesawing the rate of thought generation and the availability of cognitive control. The recurrent arrows of the Cognitive Control and Thought Generation units represent autoregressive effects. Over time, subjective reported mental states would alternate between mind wandering (MW) and focused ones (FC) (center). On the map of mental states relating to mind wandering (right, modified from Christoff et al 2016), the path of state transitions stays close to the antidiagonal. b) One of the neural activity sources reverses its effect on the cognitive control from negative to positive. The reversal changes the temporal pattern of the availability of the cognitive (center). The orange +-sign indicates the time of reversal and the red circle indicates the moment when cognitive control is still strong while NoTs is high. On the map of mental states (right), the path from FC to MW has been shifted in direction, towards the region associated with creative thinking and multitasking (CT/MT). The red circle in the leftmost panel indicates the corresponding mental state.

We considered independent effects of cognitive control and thought generation, and strived to identify their neural correlates. If one of these correlates changes the direction of its effect for a period of time, this would divert the path from the antidiagonal. Figure 3 shows an example where a neural correlate of cognitive control changes its effect from negative to positive during a shift from focused (FC) to mind wandering (MW). The change pushes the path up toward the region where multi-tasking and creative thinking are situated (MT/CT). Such an off-diagonal path has previously been reported to occur on the scale of hours in an experience sampling study (Nakatani et al., 2019). The current findings suggest that such a diverted path could occur on the scale of minutes, as a result of the direction change of neural effects on cognitive control. The change in direction of the effect might reflect a shift in mental focus from external to internal information as related studies have suggested (Andrews-Hanna et al., 2018; Buckner & Dinicola, 2019).

We proposed an approach to mental states beyond their piecemeal neural correlates. Tthe transition to mind wandering can be indirect, passing through a state that is not an intermediate on a unidimensional continuum between focused and mind wandering behavior. By situating mind wandering transitions within a multidimensional repertoire of spontaneous mental states, the current approach contributes to a unified understanding of how neural dynamics affect transitions between mental states.

## Acknowledgments

Authors would like to thank our reviewing editor Dr. Pasko Rakic and two anonymous reviewers for their encouragement and constructive advice.

For information regarding this article, please contact Chie Nakatani.

## Appendix

### 1. Mind wandering percentage (%MW) histogram

Figure A1 provides the %MW histogram over all participants. Low-, intermediate-, and high %MW trials are marked in yellow (%MW<30), gray (40≦%MW<60), and purple (%MW≧70), respectively

**Fig A1.**
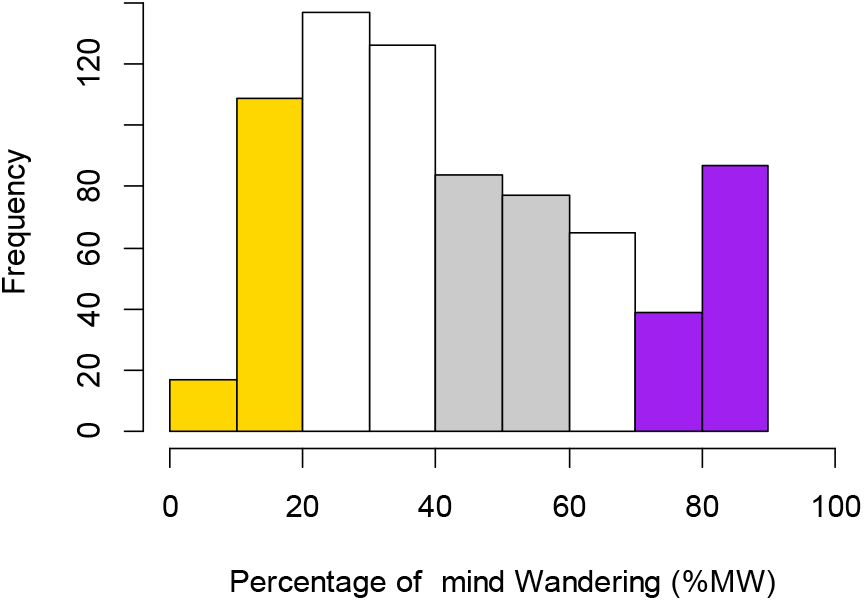
Histogram of the reported mind wandering percentage (%MW)

### 2. Electrodes for auditory evoked potential (AEP) and Alpha amplitudes

**Figure A2.**
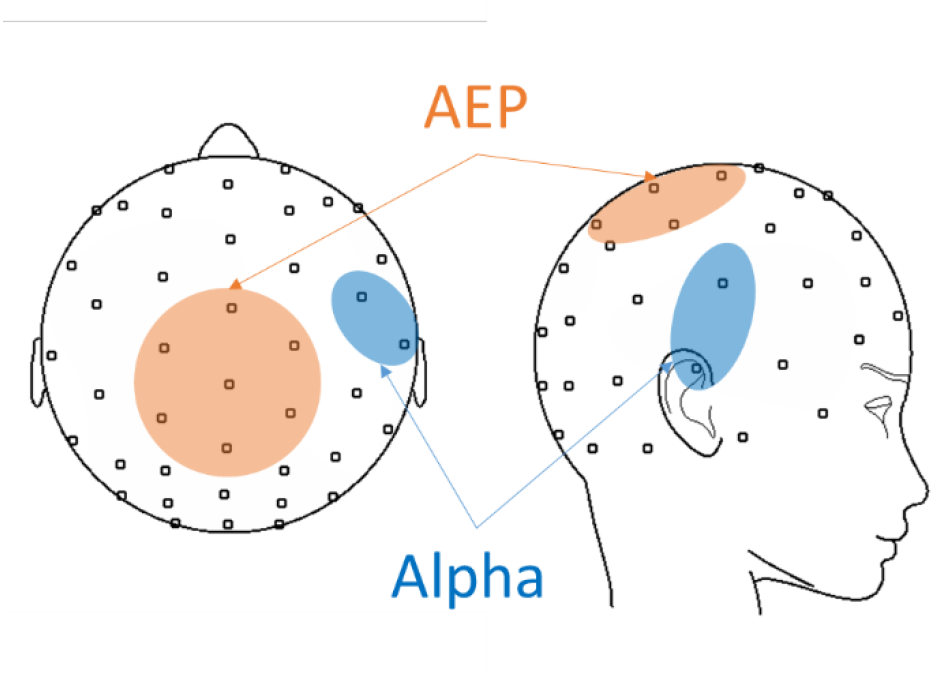
Electrodes used to compute the AEP and Alpha amplitude are indicated. Note that the electrode placement is not the international 10-10, but a manufacturer-defined system.

### 3. Alpha amplitude averaged from mid-level %MW

**Figure A3.**
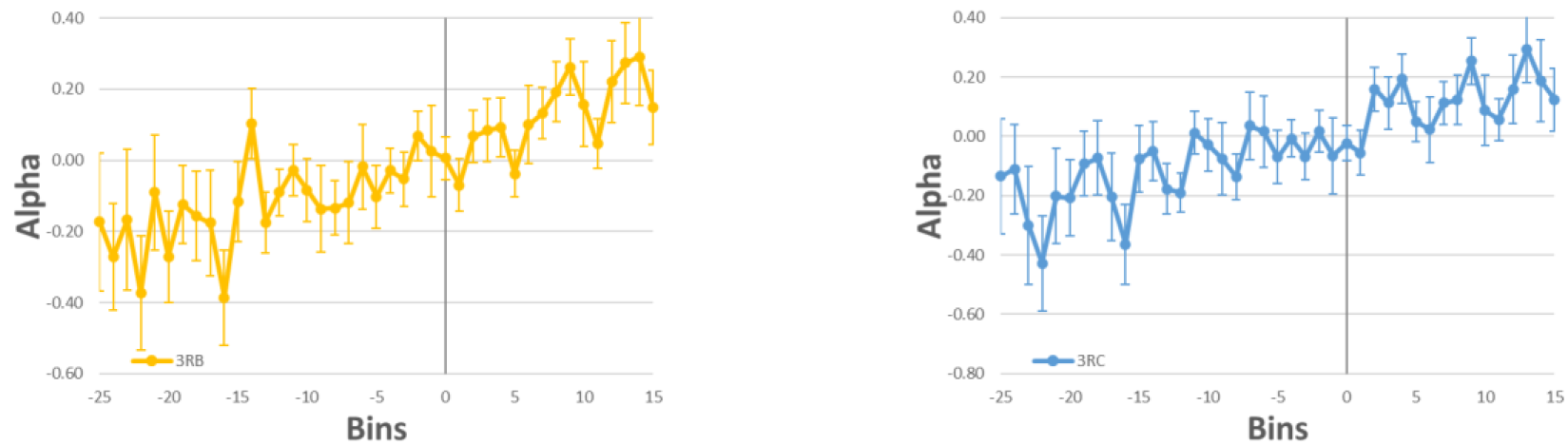
Average waveform of two right temporal electrodes, 3RB (left) and 3RC (right) Bin0 starts from the mid-level %MW report (40≦%MW<60). Error bars indicate SE.

### 4. Auto regression effects at r=1, s=1

The effect of autoregression at *i=1, j=1* was estimated to compare that with pre-selected *i\** and *l\** (See main text Notes on autoregression effect). In the same way as that in models for Tables 2, 3, and 5 in main text, the index for AEP was set as *l=1*.

a) Estimated coefficients of the full model with (*i=1, j=1, k=4, l=1*) for high %MW segments model: [*NoTs_t_*, %𝐶𝐶_t_] ∼ (*NoT*𝑠_𝑡−1_ + %𝐶𝐶_𝑡−1_) + (*Alpha*_𝑡−4_ + *AEP*_𝑡−1_) + (*Alpha*_t_ + *AEP*_t_) + *Participants*

|  | Predictors | Coefficients | Confidence interval |  | t-value | Pr(>\|t\|) |
| --- | --- | --- | --- | --- | --- | --- |
|  |  |  | 2.50% | 97.50% |  |  |
| Response var. = %CC<sub>t</sub> | (Intercept) | -4.39 | -7.07 | -1.70 | -3.21 | 0.0014 |
|  | NoTs<sub>t-1</sub> | 2.02 | -1.48 | 5.52 | 1.14 | 0.2571 |
|  | %CC<sub>t-1</sub> | 0.18 | 0.09 | 0.27 | 3.88 | 0.0001 |
|  | Alpha<sub>t-4</sub> | 8.07 | 4.11 | 12.04 | 4.00 | 0.0001 |
|  | AEP<sub>t-1</sub> | 2.46 | -3.43 | 8.35 | 0.82 | 0.4124 |
|  | Alpha<sub>t</sub> | -0.72 | -4.45 | 3.00 | -0.38 | 0.7031 |
|  | AEP<sub>t</sub> | 3.90 | -2.02 | 9.83 | 1.30 | 0.1960 |
|  | Participants | 0.36 | 0.14 | 0.59 | 3.16 | 0.0017 |
| Response var. = NoTs<sub>t</sub> | (Intercept) | 0.00 | -0.07 | 0.07 | -0.11 | 0.9120 |
|  | NoTs<sub>t-1</sub> | 0.19 | 0.10 | 0.28 | 4.06 | 0.0001 |
|  | %CC<sub>t-1</sub> | 0.00 | 0.00 | 0.01 | 2.44 | 0.0150 |
|  | Alpha<sub>t-4</sub> | 0.07 | -0.04 | 0.17 | 1.28 | 0.2020 |
|  | AEP<sub>t-1</sub> | -0.12 | -0.27 | 0.03 | -1.54 | 0.1240 |
|  | Alpha<sub>t</sub> | 0.08 | -0.02 | 0.18 | 1.64 | 0.1030 |
|  | AEP<sub>t</sub> | 0.11 | -0.04 | 0.26 | 1.41 | 0.1600 |
|  | Participants | 0.00 | 0.00 | 0.01 | 0.98 | 0.3280 |

b) Estimated coefficients of the full model with (*i=1, j=1, k=5, l=1*) for high %MW segments model: [*NoTs_t_*, %𝐶𝐶_t_] ∼ (*NoT*𝑠_𝑡−1_ + %𝐶𝐶_𝑡−1_) + (*Alpha*_𝑡−5_ + *AEP*_𝑡−1_) + (*Alpha*_t_ + *AEP*_t_) + *Participants*

|  | Predictors | Coefficients | Confidence interval |  | t-value | Pr(>\|t\|) |
| --- | --- | --- | --- | --- | --- | --- |
|  |  |  | 2.50% | 97.50% |  |  |
| Response var. = %CC<sub>t</sub> | (Intercept) | -4.50 | -7.20 | -1.80 | -3.28 | 0.0011 |
|  | NoTs<sub>t-1</sub> | 1.75 | -1.78 | 5.29 | 0.98 | 0.3293 |
|  | %CC<sub>t-1</sub> | 0.17 | 0.08 | 0.26 | 3.56 | 0.0004 |
|  | Alpha<sub>t-5</sub> | 6.74 | 2.75 | 10.72 | 3.32 | 0.0010 |
|  | AEP<sub>t-1</sub> | 3.82 | -2.09 | 9.74 | 1.27 | 0.2047 |
|  | Alpha<sub>t</sub> | 0.19 | -3.50 | 3.87 | 0.10 | 0.9212 |
|  | AEP<sub>t</sub> | 2.68 | -3.27 | 8.63 | 0.88 | 0.3769 |
|  | Participants | 0.38 | 0.15 | 0.60 | 3.28 | 0.0011 |
| Response var. = NoTs<sub>t</sub> | (Intercept) | 0.00 | -0.07 | 0.06 | -0.13 | 0.8939 |
|  | NoTs<sub>t-1</sub> | 0.18 | 0.09 | 0.28 | 3.98 | 0.0001 |
|  | %CC<sub>t-1</sub> | 0.00 | 0.00 | 0.01 | 2.30 | 0.0217 |
|  | Alpha<sub>t-5</sub> | 0.06 | -0.04 | 0.16 | 1.19 | 0.2362 |
|  | AEP<sub>t-1</sub> | -0.11 | -0.26 | 0.04 | -1.39 | 0.1656 |
|  | Alpha<sub>t</sub> | 0.09 | -0.01 | 0.18 | 1.80 | 0.0718 |
|  | AEP<sub>t</sub> | 0.10 | -0.05 | 0.25 | 1.28 | 0.2030 |
|  | Participants | 0.00 | 0.00 | 0.01 | 1.02 | 0.3088 |

c) Estimated coefficients of the full model with (*i=1, j=1, k=3, l=1*) for high %MW segments model: [*NoTs_t_*, %𝐶𝐶_t_] ∼ (*NoT*𝑠_𝑡−1_ + %𝐶𝐶_𝑡−1_) + (*Alpha*_𝑡−3_ + *AEP*_𝑡−1_) + (*Alpha*_t_ + *AEP*_t_) + *Participants*

|  | Predictors | Coefficients | Confidence interval |  | t-value | Pr(>\|t\|) |
| --- | --- | --- | --- | --- | --- | --- |
|  |  |  | 2.50% | 97.50% |  |  |
| Response var. = %CC<sub>t</sub> | (Intercept) | -6.09 | -9.89 | -2.30 | -3.16 | 0.0017 |
|  | NoTs<sub>t-1</sub> | 6.46 | 2.36 | 10.55 | 3.10 | 0.0021 |
|  | %CC<sub>t-1</sub> | 0.11 | 0.01 | 0.21 | 2.17 | 0.0310 |
|  | Alpha<sub>t-3</sub> | -10.48 | -15.12 | -5.84 | -4.44 | 0.0000 |
|  | AEP<sub>t-1</sub> | 1.53 | -4.98 | 8.04 | 0.46 | 0.6449 |
|  | Alpha<sub>t</sub> | 7.30 | 2.64 | 11.96 | 3.08 | 0.0022 |
|  | AEP<sub>t</sub> | -2.23 | -8.89 | 4.43 | -0.66 | 0.5110 |
|  | Participants | 0.52 | 0.21 | 0.83 | 3.26 | 0.0012 |
| Response var. = NoTs<sub>t</sub> | (Intercept) | -0.05 | -0.14 | 0.03 | -1.21 | 0.2285 |
|  | NoTs<sub>t-1</sub> | 0.14 | 0.04 | 0.23 | 2.88 | 0.0042 |
|  | %CC<sub>t-1</sub> | 0.00 | 0.00 | 0.00 | 2.02 | 0.0440 |
|  | Alpha<sub>t-3</sub> | 0.10 | 0.00 | 0.21 | 1.92 | 0.0558 |
|  | AEP<sub>t-1</sub> | 0.02 | -0.13 | 0.17 | 0.29 | 0.7758 |
|  | Alpha<sub>t</sub> | -0.17 | -0.27 | -0.06 | -3.13 | 0.0019 |
|  | AEP<sub>t</sub> | 0.03 | -0.12 | 0.18 | 0.37 | 0.7103 |
|  | Participants | 0.01 | 0.00 | 0.01 | 1.75 | 0.0804 |

